# Mitoxantrone inhibits and downregulates ER*α* through binding at the DBD-LBD interface

**DOI:** 10.1101/2025.01.07.631371

**Authors:** Han Wang, Yuxuan Luo, Sandeep Artham, Qianqian Wang, Yi Peng, Zixi Yun, Xinyue Li, Chen Wu, Zhenghao Liu, Kristen L. Weber-Bonk, Chun-Peng Pai, Yuan Cao, Jiangan Yue, Sunghee Park, Ruth A. Keri, Lisheng Geng, Donald P. McDonnell, Hung-Ying Kao, Sichun Yang

## Abstract

Targeting the estrogen receptor (ER or ERα) through competitive antagonists, receptor downregulators, or estrogen synthesis inhibition remains the primary therapeutic strategy for luminal breast cancer. We have identified a novel mechanism of ER inhibition by targeting the critical interface between its DNA-binding domain (DBD) and ligand-binding domain (LBD). We demonstrate that mitoxantrone (MTO), a topoisomerase II inhibitor, binds at this previously unexplored DBD-LBD interface. Using comprehensive computational, biophysical, biochemical, and cellular analyses, we show that independent of its DNA damage response activity, MTO binding induces distinct conformational changes in ER, leading to its cytoplasmic redistribution and subsequent proteasomal degradation. Notably, MTO effectively inhibits clinically relevant ER mutations (Y537S and D538G) that confer resistance to current endocrine therapies, outperforming fulvestrant in both *in vitro* and *in vivo* assays. Our findings establish domain-domain interaction targeting as a viable therapeutic strategy for ER, with translational implications for other nuclear receptors.

## Introduction

The estrogen receptor (ER or ERα) is the primary mediator of estrogen-induced mitogenic signaling in breast cancer cells^1,2^. As a nuclear hormone receptor^3^, ER contains several functional domains with particularly well-conserved regions: the DNA-binding domain (DBD) and the ligand-binding domain (LBD), which coordinate to execute hormone-induced transcriptional regulation^4,5^. Our structural studies identified a critical allosteric channel through the DBD-LBD interface that facilitates hormonal signal transmission^6^. Mutations within this interface disrupt ER transcriptional activity^6^. Similar DBD-LBD interactions have been observed across multiple members of the nuclear receptor family^7–13^, pointing to a fundamental regulatory mechanism conserved throughout this class of transcription factors.

The structural and functional understanding of the ER domain-interface prompted us to investigate its therapeutic potential. Herein we describe the identification of mitoxantrone (MTO), a topoisomerase II inhibitor^14,15^, as a small molecule that binds this interface distinct from the canonical estrogen-binding pocket. Through comprehensive analyses, we show that independent of its canonical pharmacological activity, MTO binding drives specific conformational changes in ER, leading to its subcellular redistribution and degradation. These properties enable MTO to effectively inhibit wild-type ER and variants^16–18^ that have developed resistance to current endocrine therapy, establishing the domain-domain interface as a promising target for drug discovery across the nuclear receptor family.

## Results

### MTO targets the DBD-LBD interface to inhibit ER transcriptional activity

Computational docking screens of the NIH Clinical Collection, comprising 725 compounds with established human safety profiles, identified small molecules targeting the novel ER DBD-LBD interface previously characterized by our group (**Fig. 1A**). The docking surface encompassed by eight interfacial residues (I326, Y328, W393, E397, L403, P406, N407, and L409) derived from our multidomain ER structure model^6^.

**Fig. 1.**
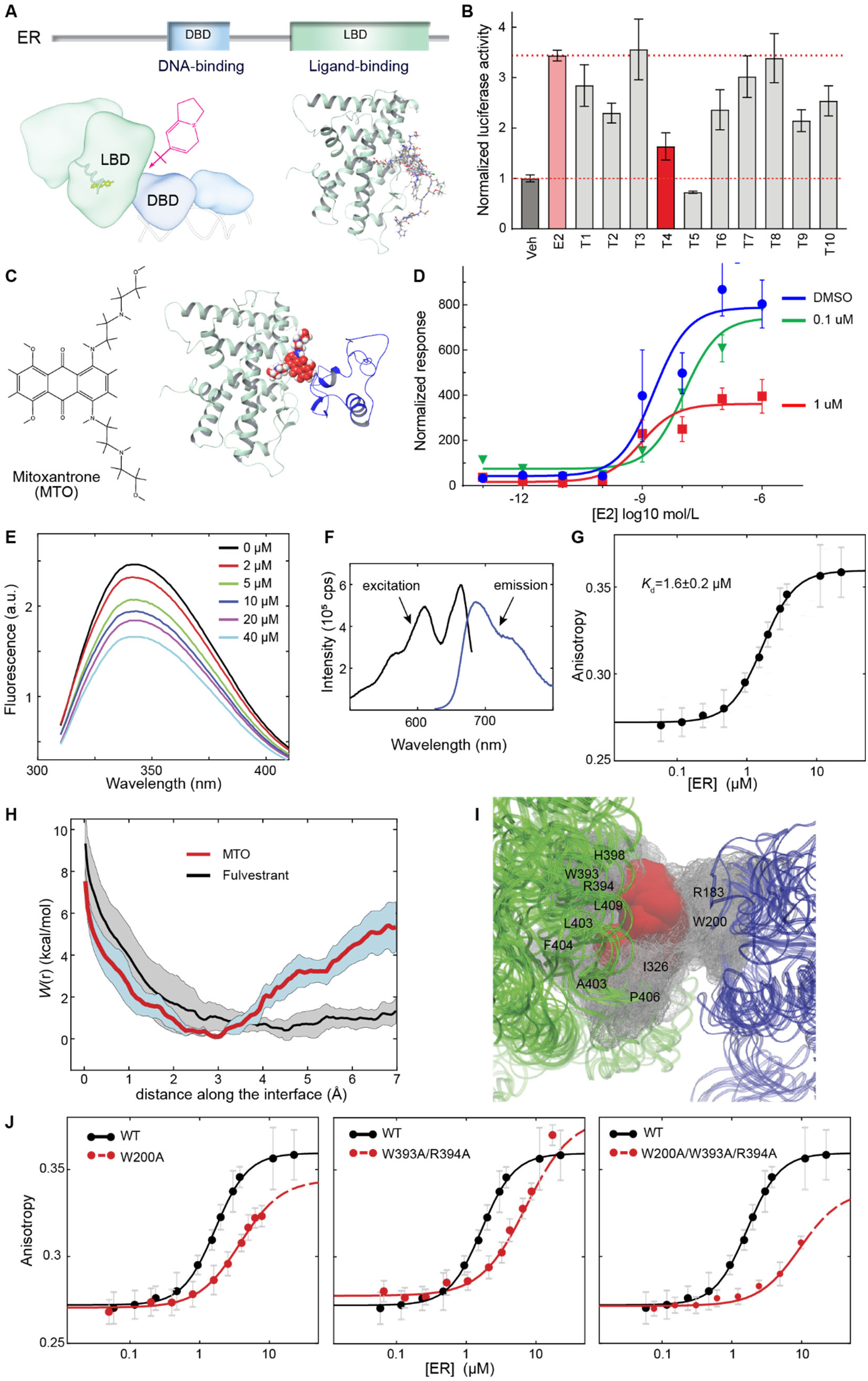
MTO binds the ER DBD-LBD interface: identification and characterization using computational, biophysical, and biochemical approaches. (**A**) Virtual docking of small molecules (pink) into the LBD-DBD interface of the multidomain ER dimer model (SASBDB access code SASDDU8)^6^. Left: ER domain organization and LBD-DBD model; right: Top 10 docked small molecules overlaid. (**B**) ERE-mediated transcription assay in HeLa cells testing top 10 candidates (1 μM, named T1-T10; Supplementary Table 1). MTO (T4) significantly reduces ER activity. (**C**) MTO structure and predicted binding pose at the DBD-LBD interface. (**D**) Dose-dependent inhibition of ER transcriptional activity by MTO in Hep-G2 cells (luciferase reporter normalized to β-galactosidase). (**E**) Tryptophan fluorescence quenching upon MTO titration demonstrates proximity to interface residue W200 (excitation at 295 nm). (**F**) MTO fluorescence characterization: excitation and emission spectra. (**G**) Fluorescence anisotropy measurements of MTO-ER binding an apparent *K*_d_ of 1.6 ± 0.2 μM. Excitation/emission: 610/685 nm. (**H**) Umbrella sampling MD simulations using MTO-ER docked model from (C). Free energy profile (solid line ± SD, shaded) along DBD-LBD interface shows specific MTO binding; fulvestrant serves as negative control. (**I**) Key amino acids at the free energy minimum. LBD (green), DBD (blue), MTO (red), interacting residues (gray, labeled). (**J**) Fluorescence anisotropy (excitation/emission, 610/685 nm) with ER mutants validates computationally predicted interactions: W200A (DBD), W393A/R394A (LBD), and W200A/W393A/R394A (triple).

Among the highest-ranked candidates (T1-T10; Supplementary Table 1), mitoxantrone (MTO) demonstrated robust inhibition of ER/E2/ERE-mediated transcriptional activity in cell-based assays (**Fig. 1B**). We included raloxifene (T5), a Selective Estrogen Receptor Modulator (SERM), as a control given its established inhibitory activity^19^. Based on its predicted binding mode at the DBD-LBD interface and therapeutic implications, we selected MTO (T4) for detailed characterization (**Fig. 1C**) (**Fig. 1C**). A cell-based luciferase ER/ERE reporter assay confirmed MTO’s dose-dependent inhibition of ER activity (**Fig. 1D**). Notably, the identified MTO binding site closely resembles the secondary tamoxifen binding site in ERβ^20,21^.

Biophysical analyses validated direct MTO binding to the DBD-LBD interface. Steady-state tryptophan fluorescence quenching showed concentration-dependent effects on the amino acid W200, positioned centrally within the DBD-LBD interface, indicating MTO’s interaction with this interfacial residue (**Fig. 1E**). Utilizing MTO’s intrinsic fluorescence properties (**Fig. 1F**), fluorescence anisotropy measurements determined an apparent binding affinity (*K*_D_) of 1.6 ± 0.2 μM (**Fig. 1G**).

Umbrella sampling molecular dynamics (MD) simulations^22^ revealed the molecular basis of this interaction. The computed potential of mean force demonstrated a favorable binding minimum corresponding to MTO stabilization at the interface (**Fig. 1H**). Analysis of protein-ligand contacts (**Fig. 1I**) demonstrated binding specificity, as the control compound fulvestrant showed no significant interface interaction (**Fig. 1H**)

Site-directed mutagenesis studies supported the predicted binding mode. Single (W200A, DBD) or double (W393A/R394A, LBD) alanine substitutions partially reduced MTO binding (**Fig. 1J**). The triple mutant (W200A/W393A/R394A), spanning both interface sides, exhibited substantially diminished binding, confirming the essential role of these residues in the MTO-receptor interaction. These complementary computational and experimental analyses establish MTO as a novel ER modulator targeting the DBD-LBD interface through specific molecular interactions.

### MTO binding induces distinct conformational changes in ER

To further characterize MTO-induced conformational changes, we employed quantitative phage-display and mammalian two-hybrid assays^23^ using a validated combinatorial peptide library that detects exposed ER surfaces^24–26^. Our analysis revealed that wildtype ER and endocrine therapy resistant-associated mutants (Y537S and D538G)^16,27–32^ adopt distinct conformational states in their unliganded apo-form, as demonstrated by their different peptide binding patterns (**Fig. 2A-C**).

**Fig. 2.**
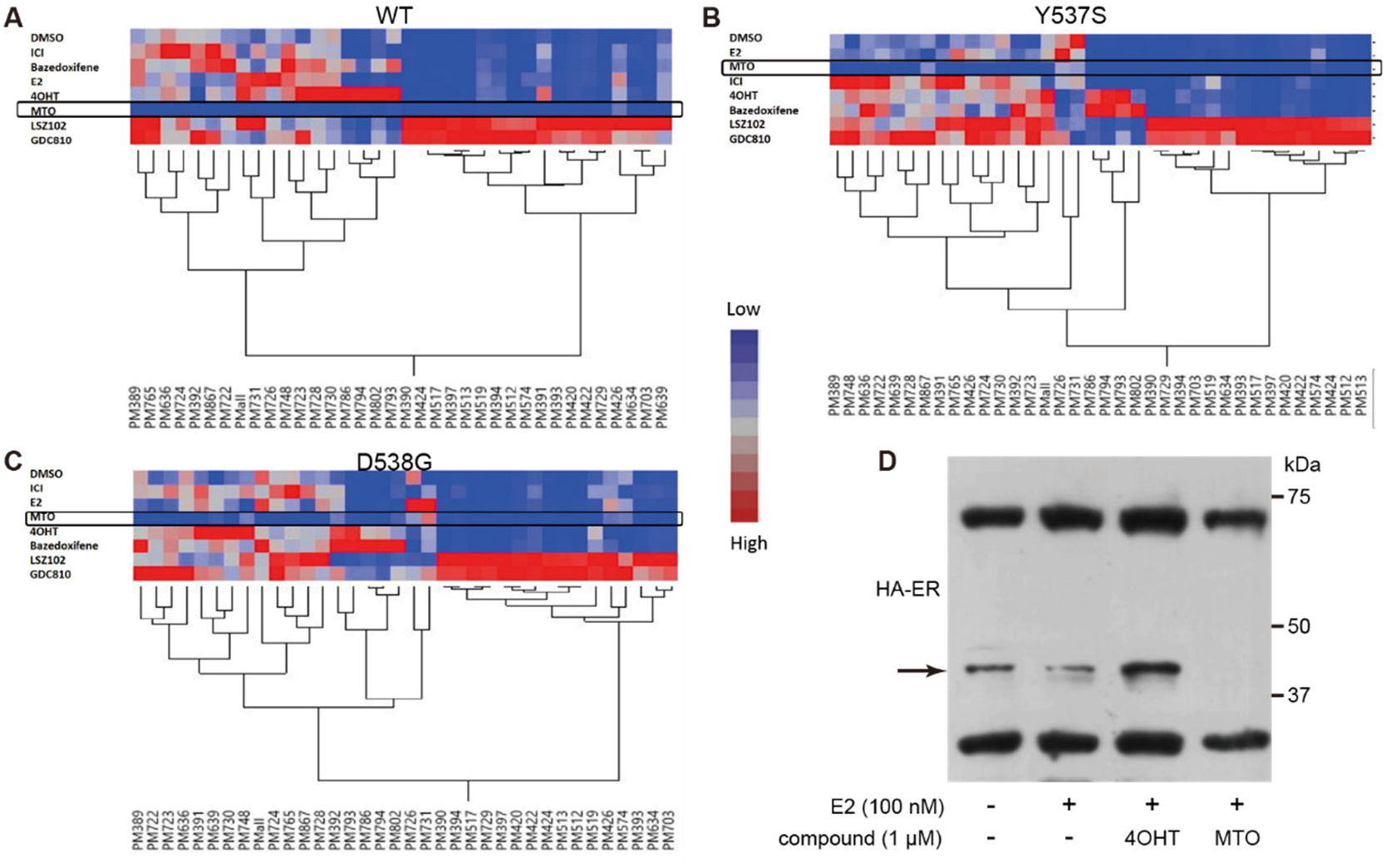
MTO alters conformational dynamics of wildtype and mutant (Y537S/D538G) ER. **(A-C)** Mammalian two-hybrid assays ER-peptide interactions in HepG2 cells. Cells were co-transfected with Gal4DBD-peptide fusions and VP16-ER constructs. Interaction strength was quantified by Gal4-responsive luciferase reporter and normalized to β-galactosidase activity. Data show MTO’s differential effects on peptide binding to (A) wild-type ER, (B) Y537S mutant, and (C) D538G mutant. Peptides were previously identified through phage display screening for ER interaction ± MTO. (**D**) Western blot analysis of HA-tagged ER proteolytic patterns in HeLa cell lysates ± MTO treatment. Arrow indicates a proteolytic fragment specifically absent after MTO treatment, suggesting ligand-induced conformational changes.

Comparative analysis with established SERMs and selective estrogen receptor downregulators (SERDs)^33,34^ (4OH-tamoxifen, bazedoxifene, fulvestrant, LSZ102, and GDC810) revealed compound-specific peptide-binding patterns, an established surrogate for conformational changes^23^. MTO treatment substantially inhibited the interactions of all peptides with wildtype ER, suggesting the receptor adopts a previously uncharacterized conformational state (**Fig. 2A**). The resistant mutants Y537S and D538G maintained selective interactions with specific peptide subsets following MTO treatment (**Fig. 2B,C**), indicating differential conformational responses between wildtype ER and disease-relevant mutations.

To quantitatively assess MTO’s impact on key protein-protein interactions, we evaluated ER-SRC3 interactions using a protein-protein interaction assay. HEK293 cells expressing ER and SRC3 were treated with vehicle (DMSO), E2, fulvestrant, or MTO. Unlike fulvestrant, which disrupted E2-mediated ER-SRC3 interaction, MTO did not affect this interaction (**Fig. S1**). This finding suggests MTO engages receptor surfaces distinct from the E2-binding pocket.

Limited proteolysis^35,36^ with optimized protease concentrations provided additional structural insights. While E2 treatment modestly reduced a specific ER fragment, MTO treatment generated a distinct proteolytic pattern characterized by complete loss of this fragment and a modest decrease in full-length HA-ER levels (**Fig. 2D**). This proteolytic signature further supports MTO-induced conformational changes. Together, these approaches demonstrate that MTO induces distinct conformational changes in wildtype and mutant ER through mechanisms different from classical SERMs and SERDs.

### MTO suppresses the phenotypes of breast cancer cells expressing disease-relevant ER mutants

MTO demonstrated superior growth inhibition compared to both 4-hydroxytamoxifen (4OH-tamoxifen) and fulvestrant in MCF7 cells harboring knock-in Y537S and D538G ER mutations^16–18^ (**Fig. 3A**). Independent validation using T47D cells with identical mutations confirmed these findings (**Fig. S2**).

**Fig. 3.**
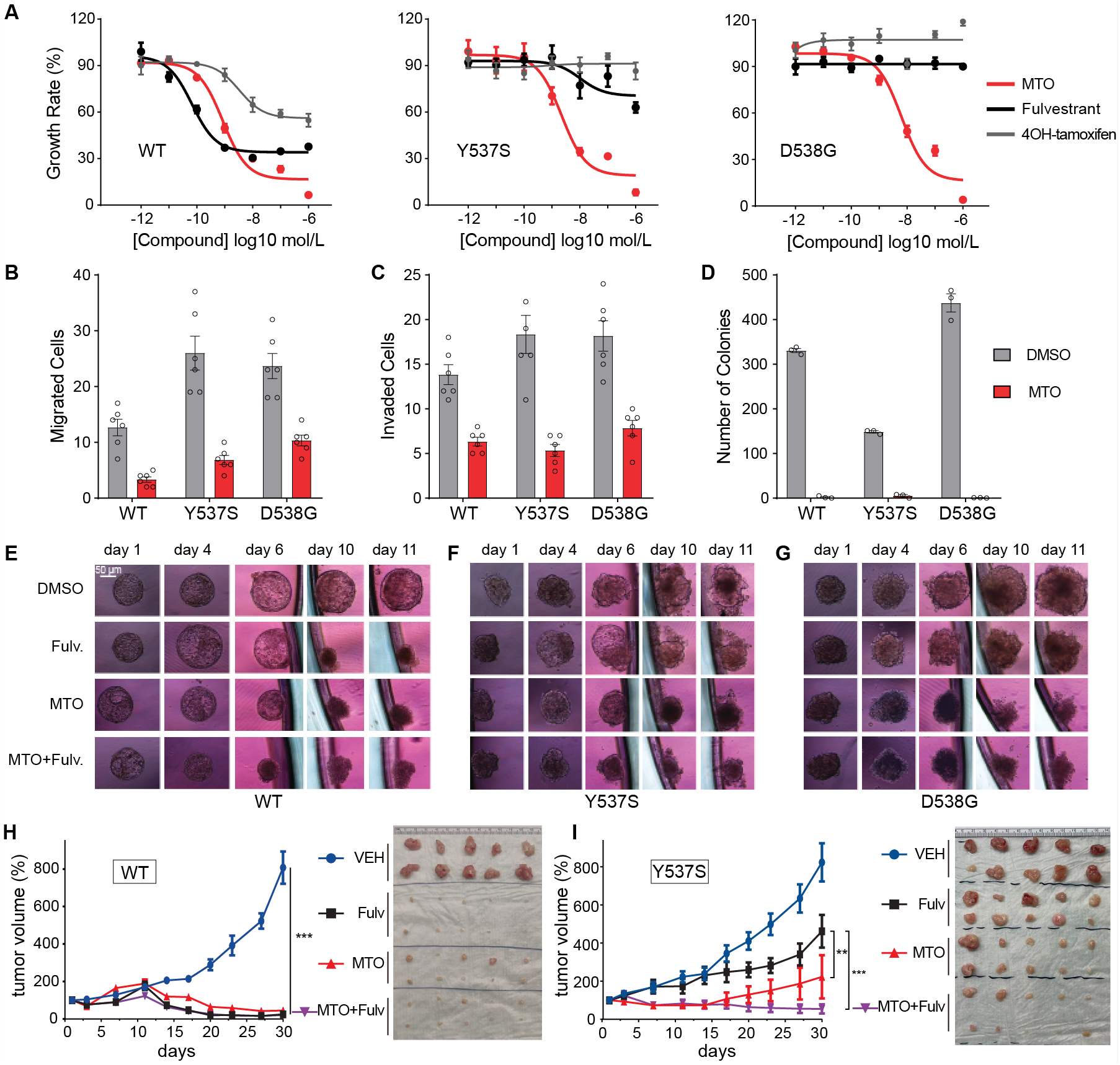
MTO demonstrates superior inhibition against ER Y537S and D538G mutant cells. (**A**) Growth inhibition analysis comparing MTO, 4OH-tamoxifen, and fulvestrant in MCF7 cells expressing knock-in ERα Y537S and D538G mutations. Cell viability was assessed using the CCK-8 assay, with IC50 values calculated for each compound. (**B**-**D**) Quantification of cancer cell phenotypes in MCF7 ER Y537S and D538G cells treated with MTO (1 μM): (**B**) cell migration, (**C**) invasion capacity, and (**D**) colony formation ability. (**E-G**) Three-dimensional spheroid growth analysis of MCF7 cells expressing (**E**) wildtype ER, (**F**) Y537S mutant, or (**G**) D538G mutant treated with MTO (1 μM) alone or in combination with fulvestrant (1 μM). Single spheroids were generated by seeding 3,000 cells in 200 μL complete growth medium and incubating for 24 hours. Treatment conditions included 0.1% DMSO (vehicle), 1 μM 4OH-tamoxifen, 1 μM fulvestrant, or 1 μM MTO. Medium was replenished with fresh treatment every 48 hours (50% volume) to maintain spheroid integrity and drug concentrations. Experiments were performed in technical triplicate and repeated independently three times. (**H**,**I**) Orthotopic xenograft studies in 6-week-old female NOD/SCID mice implanted with MCF7-WT or MCF7-Y537S cells and supplemented with estrogen pellets. Treatment was initiated when tumors reached approximately 62.5 mm³ (5 mm × 5 mm). Treatment groups (n=10 per group) received: vehicle control (VEH); fulvestrant (50 mg/kg, subcutaneous, twice weekly); MTO (1 mg/kg, intraperitoneal, weekly); or combination therapy with reduced doses of both agents (fulvestrant: 30 mg/kg, subcutaneous, twice weekly; MTO: 0.5 mg/kg, intraperitoneal, weekly). Mean ±SEM (n=10/group). One-way ANOVA with Dunnett’s test: \**p* < 0.05; \*\**p* < 0.01; \*\*\**p* < 0.005.

Treatment of ER mutant MCF7 cells with MTO (1 μM) effectively suppressed hallmark cancer phenotypes, including cell migration, invasion, and colony formation (**Fig. 3B-D**). In three-dimensional culture conditions that better approximate tumor architecture, MTO alone or combined with fulvestrant reduced growth of wildtype and mutant (Y537S, D538G) MCF7 cells more effectively than fulvestrant monotherapy (**Fig. 1E-G** and **Fig. S3**).

Comparative dose-response studies revealed more potent antiproliferative effects of MTO (0.1 and 1 μM) relative to 4OH-tamoxifen and fulvestrant (**Fig. S4**). While combinations of MTO with either agent showed comparable effects to MTO alone, the combination of low-dose MTO (0.1 μM) with fulvestrant enhanced growth inhibition. In orthotopic xenograft models, MTO demonstrated superior tumor suppression compared to fulvestrant in MCF7-Y537S tumors (**Fig. 3H,I and Fig. S5**). Notably, the combination of lower-dose MTO with fulvestrant produced substantial reductions in tumor burden.

These findings establish MTO as a potent inhibitor of endocrine therapy-resistant breast cancer cells, particularly those harboring the constitutively active Y537S and D538G ER mutations.

### MTO alters ER subcellular distribution and reduces its protein abundance

Using multiple complementary approaches, we examined the effect(s) of MTO on ER protein abundance and localization. Treatment with MTO (1 μM) decreased ER protein levels in a time-dependent manner in both MCF7 and T47D cells expressing wildtype or Y537S/D538G knock-in mutant ER (**Fig. 4A,B**). This reduction affected both endogenous ER and exogenously expressed HA-tagged Y537S mutants in MCF7 cells (**Fig. 4C**). Immunofluorescence microscopy and quantitative in-cell western assays confirmed dose-dependent ER reduction in MCF7 cells treated with MTO (0.1-1 μM, 48 hours), normalized to DNA content (**Fig. 4D,E**).

**Fig. 4.**
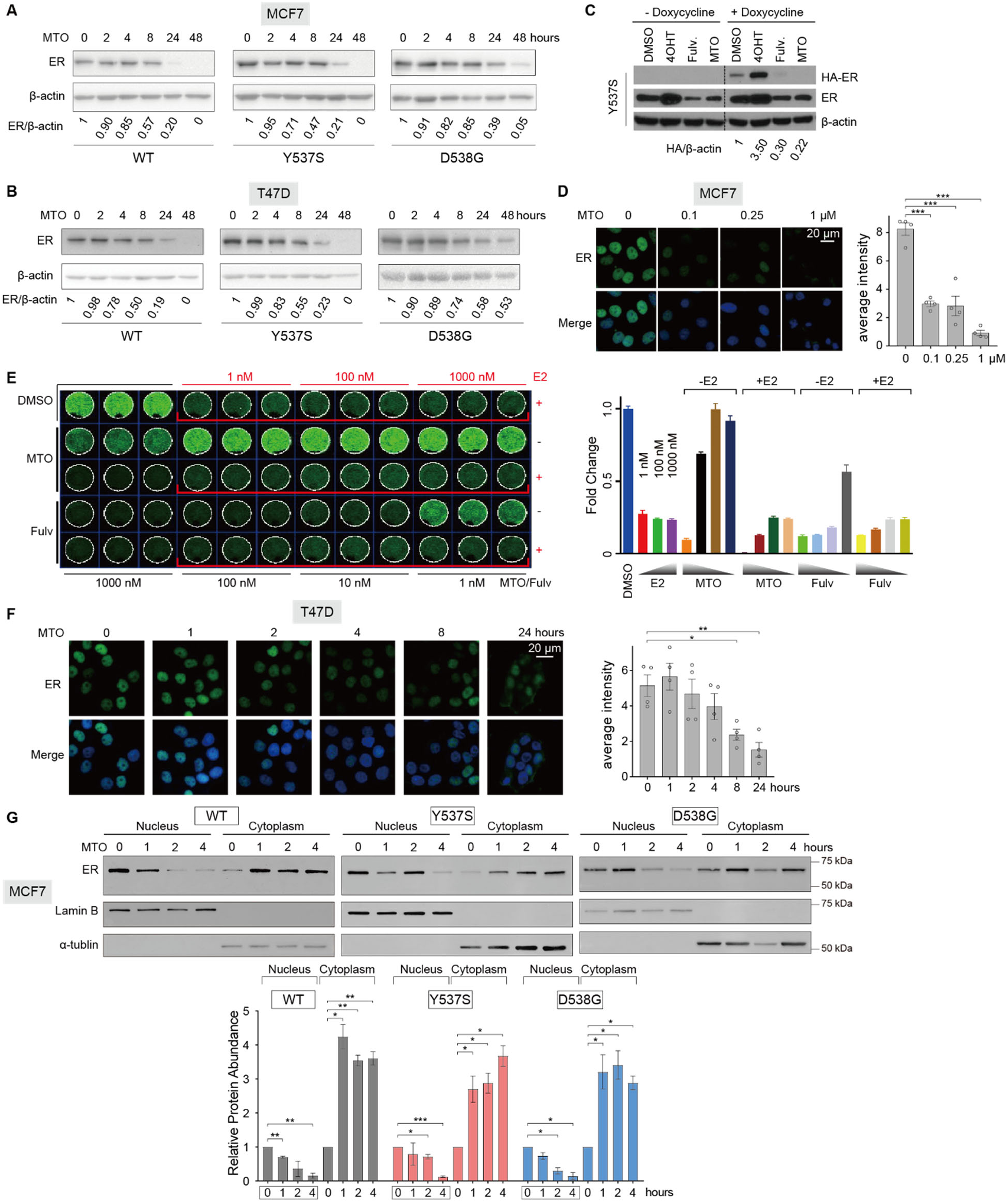
MTO reduces ER abundance and alters its subcellular distribution. (**A,B**) MTO (1 μM) treatment reduces ER protein levels over time in MCF7 (**A**) and T47D (**B**) cells expressing wildtype or Y537S/D538G knock-in mutant ER, quantified relative to β-actin. (**C**) In MCF7 cells expressing exogenous HA-tagged Y537S ER mutants, MTO decreases both endogenous and exogenous ER levels, with HA-tagged mutant quantified relative to β-actin. (**D**) Immunofluorescence microscopy reveals dose-dependent reduction of ER protein in MCF7 cells treated with MTO (0.1, 0.5, and 1 μM, 48 hours). (**E**) In-cell western assay shows ER levels in MCF7 cells after 18-hour treatment with E2, MTO, or fulvestrant, normalized to DNA content (DRAQ5) and expressed as percentage of DMSO control. (**F**) Time-course immunofluorescence in T47D cells shows nuclear ER reduction after 2-hour MTO treatment. (**G**) Subcellular fractionation reveals early (2-hour) ER accumulation in cytoplasm with concurrent nuclear depletion. Data represent mean ±SEM (n=4). One-way ANOVA with Dunnett’s test: \**p* < 0.05; \*\**p* < 0.01; \*\*\**p* < 0.005.

Detailed analysis of early time points revealed a distinct sequence of events. While total ER levels remained stable at 2 hours post-treatment (**Fig. 4B**), immunofluorescence showed substantial nuclear depletion (**Fig. 4F**). Nucleocytoplasmic fractionation studies confirmed a rapid decrease in nuclear ER within 1 hour of MTO treatment in wildtype cells, accompanied by modest changes in cytosolic ER abundance through 4 hours (**Fig. 4G**). The Y537S and D538G mutants exhibited similar redistribution patterns but with delayed nuclear depletion at 2 hours post-treatment, aligning with immunofluorescence observations (**Fig. 4B**).

These findings demonstrate that MTO acts through a distinct mechanism: first inducing ER redistribution from the nucleus to the cytoplasm, followed by time-dependent protein degradation. This pathway contrasts with classical SERDs such as fulvestrant, which promote ER sequestration in an insoluble nuclear fraction before proteasomal degradation^5,37^.

### MTO and fulvestrant induce distinct transcriptional responses

To elucidate the mechanism by which MTO impacts ER-mediated transcription, we performed RNA-seq analysis on MCF-7 cells expressing wildtype ER or Y537S/D538G mutants treated with vehicle, MTO, fulvestrant, or with both drugs. Western blot analysis confirmed differential effects on ER protein levels across treatments and cell lines after 48 hours (**Fig. 5A**). Principal component analysis of global gene expression revealed distinct clustering by treatment and cell line, with the first two principal components (PC1 and PC2) accounting for 45% of the total variance and primarily distinguishing the cell lines (**Fig. 5B**).

**Fig. 5.**
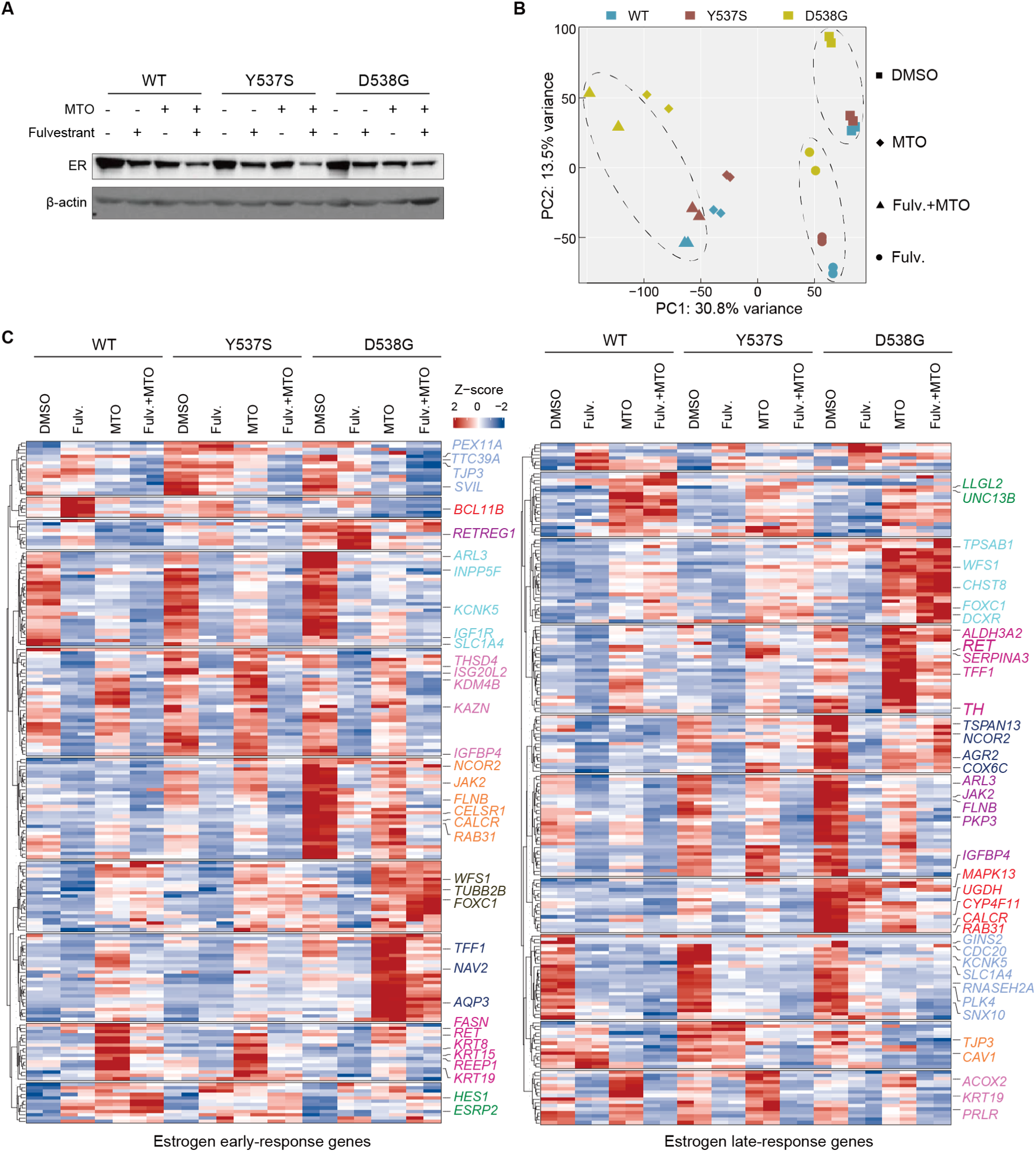
MTO distinctly regulates ER target gene expression compared to fulvestrant. (**A**) Western blot analysis of ER protein levels in wildtype, Y537S, and D538G knock-in MCF7 cells treated for 48 hours with vehicle, fulvestrant (1 μM), MTO (1 μM), or combination. β-actin serves as loading control. (**B**) Principal component analysis of global gene expression profiles in wildtype, Y537S, and D538G knock-in MCF7 cells treated for 48 hours with vehicle, fulvestrant (1 μM), MTO (1 μM), or combination. Each point represents a distinct treated cell line. PC1 and PC2 explain 31% and 14% of total variance, respectively. (**C**) Heatmaps showing early (4-6 hours) and late (24 hours) estrogen-responsive gene expression in wildtype, Y537S, and D538G knock-in MCF7 cells. Z-score normalized expression values: red indicates upregulation, blue indicates downregulation versus vehicle control. Highlighted genes represent ER-bound promoters. Soft clustering performed using Mfuzz package in *R* with fuzzy c-means algorithm.

Differential gene expression analysis through hierarchical clustering revealed compound-specific effects on established early (4-6 hours) and late (24 hours) estrogen-responsive genes^38^ across the isogenic cell panel (**Fig. 5C**). MTO significantly modulated DNA-damage-response pathways, consistent with its known mechanisms of action, while fulvestrant demonstrated cell line-dependent transcriptional effects (**Fig. S6A**). Gene Set Enrichment Analysis identified both overlapping and distinct effects on cellular signaling pathways between the two compounds (**Fig. S6B**).

Analysis of oncogenic signatures revealed compound-specific regulatory patterns, with MTO selectively modulating specific transcriptional programs while fulvestrant exhibited broader effects on gene expression (**Fig. S7A**). Notably, in endocrine therapy-resistant Y537S and D538G mutants, MTO demonstrated more pronounced effects on proliferation-associated signatures compared to fulvestrant (**Fig. S7B,C**).

### MTO promotes proteasome-dependent ER degradation

After establishing that MTO induces distinct conformational changes in ER that correspond with enhanced protein turnover, we sought to elucidate the molecular mechanisms underlying this downregulation. Protein turnover analyses were conducted using MCF7 and T47D cells expressing either wildtype ER or Y537S/D538G knock-in mutations. Addition of cycloheximide to block new protein synthesis demonstrated that MTO treatment substantially decreased ER protein half-life across all cell variants (**Fig. 6A,B**).

**Fig. 6.**
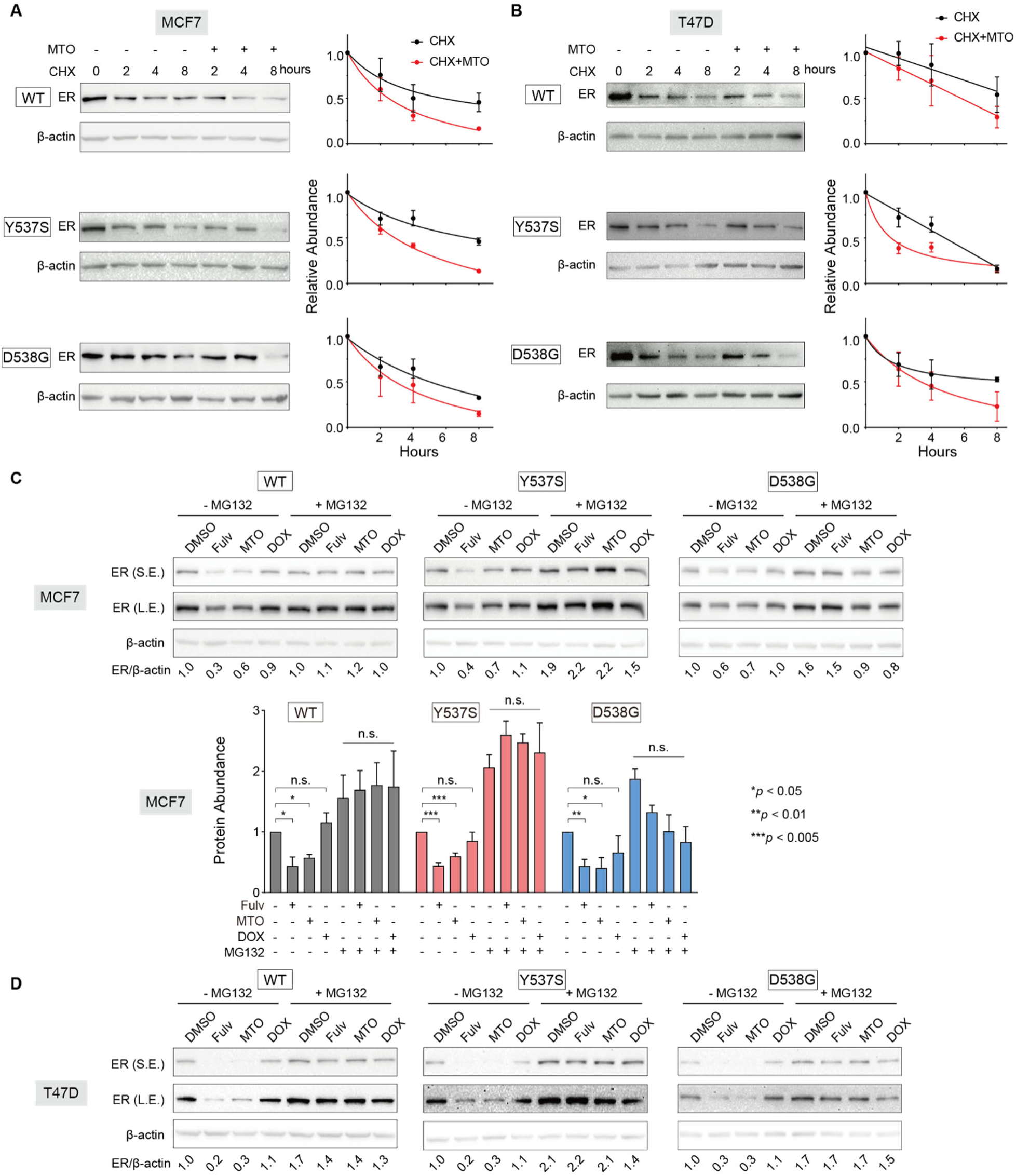
MTO promotes proteasome-dependent ER degradation. (**A**,**B**) Protein turnover assays in MCF7 (**A**) and T47D (**B**) cells expressing wildtype, Y537S, or D538G knock-in ER. Cells were pre-treated with cycloheximide (CHX, 30 minutes) followed by DMSO or MTO (1 μM) for 0-8 hours. Western blot analysis shows ER levels normalized to β-actin, with 0-hour timepoint set as 1. MTO reduces ER protein half-life compared to controls (**C**,**D**) Proteasome inhibition studies in MCF7 (**C**) and T47D (**D**) cells. Following MG132 pre-treatment (20 μM, 30 minutes), cells were treated with DMSO, fulvestrant (1 μM), MTO (1 μM), or doxorubicin (DOX, 1 μM) for 8 hours. Western blot analysis shows ER levels normalized to β-actin (n=3 biological replicates for MCF7). MG132 blocks both fulvestrant- and MTO-induced ER degradation, while structural analog doxorubicin shows little or no effect on ER stability. One-way ANOVA with Dunnett’s test: \**p* < 0.05; \*\**p* < 0.01; \*\*\**p* < 0.005.

To identify the specific degradation pathway, we investigated proteasomal involvement through pretreatment with the proteasome inhibitor MG132, followed by exposure to DMSO, fulvestrant, MTO, or doxorubicin (DOX). Notably, proteasome inhibition prevented MTO-induced ER degradation in both parental and mutant cell lines (**Fig. 6C,D**), establishing that MTO mediates ER degradation through a proteasome-dependent pathway

### Ectopically expressed ER partially rescues MTO’s antiproliferative effects

We investigated whether MTO’s growth inhibitory activity depends on its ER downregulation mechanism by examining how increased ER expression affects MTO’s antiproliferative effects. MCF7 cells expressing wildtype ER or Y537S/D538G knock-in mutations were transfected with either an ER expression vector or control vector, then treated with vehicle or MTO (0.1, 0.2 μM). Cell viability measurements using CCK-8 assays at day 5 showed that increased ERα expression partially reversed MTO’s growth inhibitory effects across all cell variants (**Fig. 7A**), suggesting ER degradation contributes to but does not fully account for MTO’s antiproliferative effects.

**Fig. 7.**
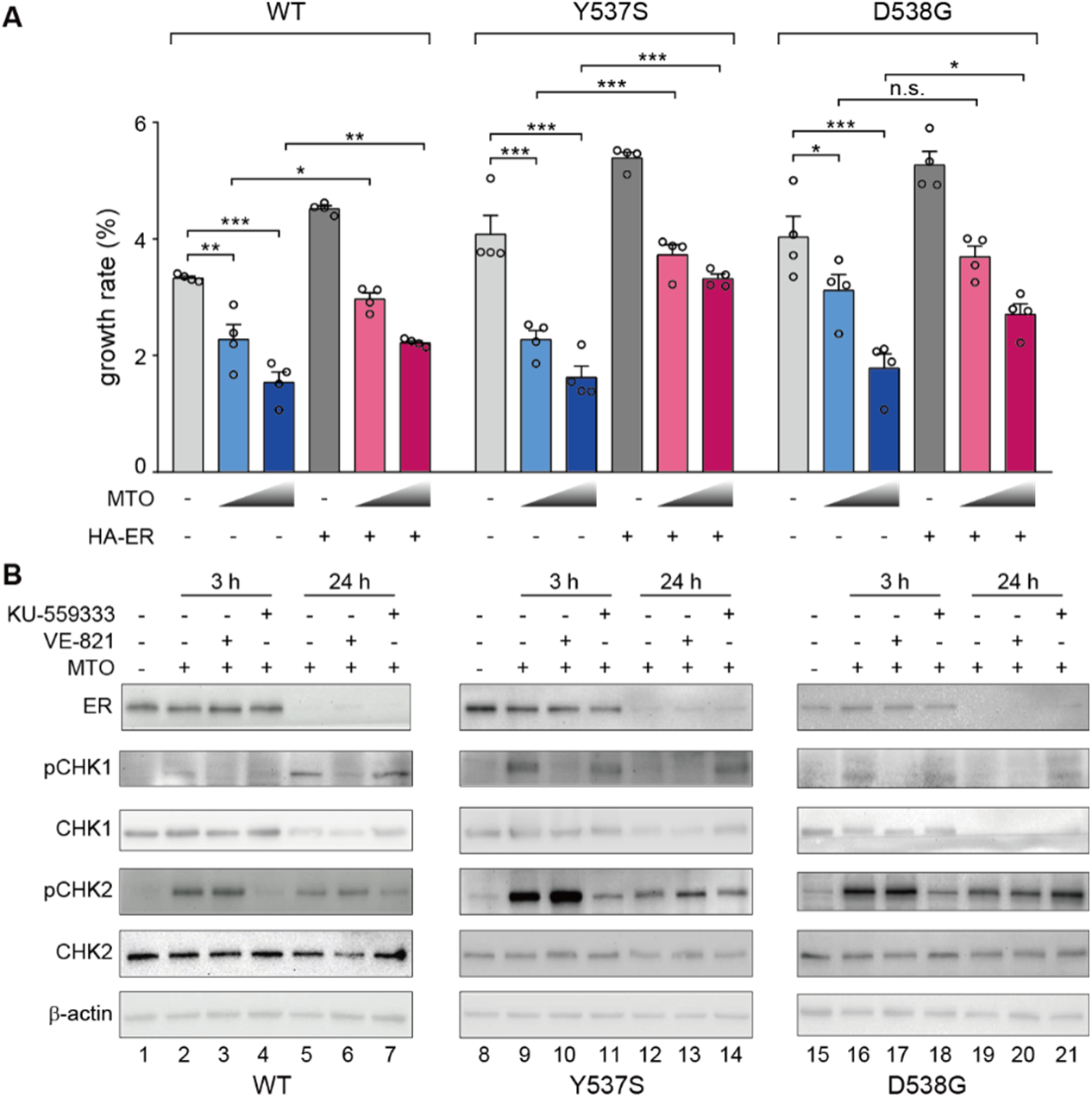
MTO-induced ER degradation occurs independently of DNA damage and affects cell survival. (**A**) Cell viability of wildtype, Y537S, and D538G knock-in MCF7 cells transfected with control or ER-expressing vector, treated with vehicle or MTO (0.1, 0.2 μM). CCK-8 assay at day 5, normalized to day 1. Mean ±SEM (n=4). One-way ANOVA with Dunnett’s test: \**p* < 0.05; \*\**p* < 0.01; \*\*\**p* < 0.005. (**B**) Western blot analysis of wildtype, Y537S, and D538G knock-in MCF7 cells pre-treated with vehicle, ATM inhibitor KU-55933 (10 μM), or ATR inhibitor VE-821 (5 μM) for 30 minutes before MTO (1 μM) treatment. ER levels and DNA damage response assessed by phospho-Chk1(S345) and phospho-Chk2(T68) as ATR and ATM readouts.

### MTO reduces ER independently of DNA-damage response

TO evaluate whether MTO’s established DNA damage response (DDR) pathway through topoisomerase IIα (Top2α) and ATM/ATR kinases^39,40^ contributes to ER downregulation, we examined the effects of specific kinase inhibitors. We pretreated wildtype and Y537S/D538G knock-in MCF7 cells with either the ATM inhibitor KU-55933 or the ATR inhibitor VE-821 prior to MTO exposure. Western blot analysis revealed that neither inhibitor prevented MTO-induced ERα reduction, despite effectively blocking ATM/ATR activation as evidenced by decreased Chk2(T68) and Chk1(S345) phosphorylation (**Fig. 7B**). These results suggest that MTO reduces ERα through mechanisms distinct from Top2α-mediated DNA damage response.

## Discussion

### The DBD-LBD interface: an emerging therapeutic target

Therapeutic strategies for targeting the ER have historically centered on the LBD^5,41,42^. While this approach has proven to be an effective strategy for developing drugs in the frontline, even contemporary LBD-targeting agents show limited efficacy in advanced disease settings^16,17^. Our study reveals the interface between the DBD and LBD of ER as a promising therapeutic target. Previous structural studies from our group established this interface as a critical allosteric channel mediating signal transmission from hormone binding to DNA binding^6^. The newly characterized binding site at this DBD-LBD interface, while distinct from the canonical estrogen-binding pocket, shares important structural features with the secondary tamoxifen binding site identified in ERβ^20,21^. These findings demonstrate the therapeutic potential of targeting domain-domain interactions in ERα and suggest valuable opportunities for drug development beyond the traditional LBD-focused approach.

### MTO’s novel mechanism of action and therapeutic implications

Our research reveals that MTO operates through a unique mechanism targeting the ER DBD-LBD interface, distinguishing it from conventional antagonists such as fulvestrant. MTO exhibits a distinctive two-phase action: first inducing rapid cytoplasmic redistribution of ER within two hours, followed by proteasome-dependent degradation.

In comprehensive testing across cellular and animal models, MTO demonstrates marked superiority over fulvestrant in treating endocrine therapy-resistant breast cancer cells, particularly those harboring Y537S and D538G ER mutations. This enhanced efficacy stems from MTO’s dual mechanism: it simultaneously inhibits ER activity and promotes ER degradation, contrasting with fulvestrant primary reliance on antagonistic effects rather than its SERD properties^39^. Other emerging compounds like ErSO achieve anti-cancer effects through different pathways such as triggering the unfolded protein response and subsequent necrosis^43^, although MTO’s complete mechanism remains under investigation.

The ability to overcome mutations that confer resistance to conventional therapies has significant therapeutic implications. Our mechanistic studies provide a molecular framework for understanding how MTO affects ER stability and function. As new ER-targeting agents emerge, such as imlunestrant^44^ and elacestrant^45,46^, resistance to endocrine therapies remains a critical challenge, particularly for ER mutants^47^. While MTO may have limited clinical utility, elucidating its mechanism of action provides valuable insights for developing therapeutic strategies beyond traditional LBD-targeting approaches.

### Translational implications of targeting DBD-LBD interfaces in nuclear receptors

Our studies establish the targeting of domain-domain interfaces as a viable therapeutic strategy for nuclear receptors. The DBD-LBD interface we characterized in ER represents a conserved regulatory mechanism found throughout the nuclear receptor family^7–13^. Identifying similar interactions in other nuclear receptors, including the glucocorticoid receptor (GR) and retinoid X receptor (RXR), points to the broad applicability of this therapeutic approach.

The androgen receptor (AR) presents a compelling example of this conservation. Two independent research groups have documented various modes of DBD-LBD interactions in AR^10,48^. While these studies were conducted at moderate resolution (9-11 Å and 13 Å), they established the fundamental importance of these domain interactions in nuclear receptor function. Despite growing documentation of DBD-LBD interactions across the nuclear receptor family, their potential as therapeutic targets has remained largely unexplored.

Our proof-of-concept study demonstrates that targeting the ER DBD-LBD interface can yield effective translational outcomes. This novel approach extends beyond traditional ligand-binding pocket strategies and suggests new therapeutics in diseases where aberrant nuclear receptor signaling drives pathology.

### Limitations and future directions

A central challenge in our study is the difficulty of definitively separating MTO’s effects on ER from its established DNA damage response (DDR) pathway. Although our evidence indicates that MTO acts via a distinct mechanism at the DBD-LBD interface, complete separation of these activities remains a key limitation. We have demonstrated that MTO’s effects on ER persist even when DNA damage response pathways are blocked through ATM/ATR kinase inhibition, and we have observed a specific temporal sequence of ER-related events. Additionally, our finding that doxorubicin, a structurally similar DNA-damaging compound, does not affect ER levels provides important evidence for mechanism specificity. However, these observations, while supportive, cannot entirely resolve the limitation.

Future research efforts to advance the therapeutic potential of targeting the DBD-LBD interface should address two critical priorities. First, high-resolution structural characterization of MTO binding will be essential for understanding the precise molecular interactions at this interface. Second, developing optimized MTO derivatives with enhanced specificity for the DBD-LBD interface could minimize off-target effects related to DNA damage while maintaining efficacy against resistant ER mutations.

## Methods and Materials

### Molecular docking

In silico docking was performed to investigate small molecule binding to the LBD-DBD interface using the Schrodinger package^49^ (version 2024-1). The LBD surface was defined based on eight interfacial residues (I326, Y328, W393, E397, L403, P406, N407, and L409) identified from a multidomain ER structure model^6^.

Virtual screening was conducted using the NIH Clinical Collection (NCC), a curated library of 725 small-molecule compounds with documented safety profiles in human clinical trials. The NCC library was procured from the National Center for Advancing Translational Sciences (Bethesda, MD) through Evotec. The compounds were supplied as 10-mM stock solutions in dimethyl sulfoxide (DMSO) solvent, facilitating direct validation of in silico predictions.

### Transient transfection reporter assay

Cells were seeded at a density of 1×10^6^ per 6 cm plate in phenol red-free DMEM supplemented with charcoal-stripped FBS. The seeding was performed one day prior to transfection. Transfections were performed using Lipofectamine 3000 (Thermo Fisher, #L3000001) with 2 μg ERE-TK-Luciferase reporter plasmid (Addgene, #11354) and 0.2 μg pRL-TK Renilla Luciferase plasmid (Addgene, #27163) per plate. Cells were cultured in charcoal-stripped serum for hormone-dependent assays for 48 hours before transfection. Post-transfection (24 hours), cells were treated with either DMSO vehicle (Thermo Scientific, #T038181000) or 100 nM 17β-estradiol (E2; Millipore Sigma, #3301) for 24 hours. Luciferase activities were measured using the dual-Luciferase reporter assay system (Promega, #E1910) on a Varioskan LUX multimode microplate reader (Thermo Fisher, #VL0000D0). Firefly luciferase activity was normalized to Renilla luciferase activity. All experiments were performed in triplicate.

In separate experiments, HeLa (ATCC, #CRM-CCL-2) or HEK293 (ATCC, #CRL-3216) cells were transiently transfected with an ERE-containing reporter construct (Addgene, #11354), an HA-tagged ER expression plasmid^50^, and a pCMV β-galactosidase plasmid (Addgene, #27163). At 24 hours post-transfection, cells were treated ± 100 nM E2, ± 1 μM 4-hydroxytamoxifen (4OH-tamoxifen; Cayman, #17308), or ± 1 μM MTO (ApexBio, #B2114) for 48 hours. Luciferase activity was normalized to β-galactosidase activity. Parallel experiments with HA-tagged ER-transfected HeLa cells underwent identical treatments, followed by Western blot analysis using anti-HA antibody (Santa Cruz, #sc-7392, 1:4000 dilution).

### Dose-dependent transcriptional activity

HepG2 cells were co-transfected with three plasmids: a 3×ERE-TATA-Luc reporter plasmid, an RST7-ERα expression plasmid, and a pCMV-β-gal plasmid. After 24 hours, cells were treated with various concentrations of E2, with or without MTO, for 48 hours. Luciferase activity was measured and normalized to β-galactosidase activity. Dose-response analysis was performed using GraphPad Prism V8 (GraphPad Software, San Diego, CA, USA).

### Recombinant protein expression and purification

The human ER segment (amino acids E181–P552), containing the DBD and LBD, was recombinantly expressed in *E. coli* BL21(DE3) cells using the pMCSG7 vector in the presence of 10 μM E2, as previously described^6^. A 19-bp double-stranded DNA duplex (ERE-DNA) containing the consensus estrogen response element was formed by annealing complementary oligonucleotides (5’-tAGGTCAcagTGACCTgcg-3’ and 5’-cgcAGGTCActgTGACCTa-3’), mirroring the sequence used in the DBD crystal structure (PDB entry 1HCQ^51^). Purified ER proteins were incubated with ERE-DNA and TIF2 peptide (KENALLRYLLDKDD) at molar ratios of 1:1.2 and 1:3.0, respectively. ER-DNA complexes were isolated by size exclusion chromatography using a Superdex 200 column in a buffer of 10 mM CHES (pH 9.5), 125 mM NaCl, 5 mM KCl, 4 mM MgCl2, 50 mM arginine, 50 mM glutamate, 5 mM TCEP, 5% glycerol, 10 µM zinc acetate, and 10 µM E2. ER mutants were generated by site-directed mutagenesis and purified following the same protocol.

### Intrinsic tryptophan fluorescence quenching

An ER-specific tryptophan fluorescence assay was previously engineered to monitor the microenvironment of Trp200 (W200), located at the LBD-DBD interface. Four native tryptophan residues were site-specifically mutated to phenylalanine, leaving W200 as a single fluorescent probe^6^. Emission spectra were recorded using a FluoroMax-4/Plus spectrofluorometer (Horiba Scientific) with an excitation wavelength of 295 nm and 2 nm bandpass. Measurements were conducted at 25 ± 0.1°C using samples containing 0.1 mg/ml ER complex in the absence or presence of MTO. Each spectrum was buffer-corrected and represents the average of three independent measurements.

### Fluorescence anisotropy

To determine the optimal excitation and emission wavelengths for mitoxantrone (MTO), its excitation and emission spectra were initially measured using a FluoroMax-4/Plus spectrofluorometer (Horiba Scientific) with a 1 nm bandpass. Based on these spectra, fluorescence anisotropy assays were performed to measure the anisotropy of 0.5 μM MTO in the absence or presence of various ER concentrations. Each measurement used an automated FL-1044 dual polarizer (FM4-Pol) with excitation and emission wavelengths set to 610 and 685 nm, respectively, and a 5 nm bandpass for both. Temperature was maintained at 25 ± 0.1°C throughout the experiments. Anisotropy values were calculated from the average of 10 replicate measurements, with standard deviations reported.

### Umbrella sampling molecular dynamics simulations

System preparation: Crystal structures of mitoxantrone (PDB entry 4G0V^52^) and fulvestrant (PDB entry 4J03^53^) were used as ligands, while the receptor was based on a multidomain ER structure model^6^. Receptor-ligand complexes were generated by initially positioning ligands near the domain interface. Complexes were solvated in a 155 Å cubic box with TIP3P water, 150 mM NaCl, and counterions for charge neutrality.

Simulation parameters: Simulations were performed using NAMD 2.13^54^ with the CHARMM22 force field^55^. Electrostatic interactions were treated using particle-mesh Ewald (14 Å cutoff for non-bonded interactions). Van der Waals interactions were switched to zero between 10-12 Å. A 2-fs time step was used with Langevin dynamics at 301 K and 1 atm. Equilibrium simulations (10 ns) were conducted with RMSD restraints on heavy atoms (100 kcal/molꞏÅ² for backbone, 10 kcal/molꞏÅ² for sidechain) to prepare systems for umbrella sampling.

Steered molecular dynamics (SMD): SMD generated 80 initial poses by pulling the ligand along the domain interface (constant velocity: 2.5×10^-3^ Å/ps). The center-of-mass distance (D) between the ligand and key interfacial residues (Y195, H196, V199, W200, W393, L403, P406, L409, E444, and F445) was monitored. Poses were spaced at 0.125 Å intervals (D: 0-10 Å). SMD simulations ran for 12 ns (mitoxantrone) and 15 ns (fulvestrant). Umbrella sampling: The ligand was harmonically restrained to its target distance (force constant: 100 kcal/molꞏÅ^2^). The ER protein was equilibrated by applying RMSD restraints on heavy atoms: 100 kcal/molꞏÅ^2^ (backbone) and 10 kcal/molꞏÅ^2^ (sidechain) for individual DBDs and LBDs, and 50 kcal/molꞏÅ^2^ (backbone) for the entire DBD-LBD complex. Each of the 80 umbrella trajectories ran for 3 ns, resulting in a cumulative simulation time of 240 ns.

Data analysis: The binding free energy profile was calculated using the weighted histogram analysis method (WHAM)^56^. Uncertainty was evaluated via block-averaging^57^. Trajectories were split into ten 300 ps blocks after 150 ps equilibration (0.1 Å bin, 1×10^-4^ tolerance). Ten receptor-ligand structures near the binding free energy minimum (D ≈ 3 Å, range: 2.2-3.5 Å) were randomly selected for additional 50 ns simulations, with ER protein RMSD-restrained and ligand unrestrained.

### Phage display for identifying ligand-specific conformational probes

Phage display experiments were conducted to identify peptides serving as conformational probes for various ER-ligand complexes. Purified ER (4 pmol) was immobilized on 96-well plates coated with consensus estrogen response element (ERE) DNA, mimicking the receptor’s native conformation. Three phage libraries (LXXLL [19 amino acids], CoRNR [23 amino acids], and X13 [13 amino acids]), each containing 10^10^ unique peptides, were incubated with immobilized ER in the presence of specific ligands for 3 hours at 25°C^23,24^. After washing to remove unbound phage, bound phage was eluted and amplified in E. coli DH5αF′. This process was repeated for five rounds of selection for each ER-ligand complex. Phage ELISA quantified enrichment of selectively bound phages. Enriched phage pools were subcloned into a mammalian two-hybrid (M2H) vector for DNA sequencing and characterization of receptor binding profiles in M2H assays.

### Mammalian two-hybrid (M2H) assay

Representative peptides from each identified class were fused to the Gal4-DBD. HepG2 cells were transiently co-transfected with four plasmids: ERα-VP16 fusion, peptide-Gal4 DBD fusion, luciferase reporter driven by five Gal4 response elements, and CMV-β-galactosidase for transfection normalization. Transfected cells were treated with various ligands (1 μM) for 24 hours in hormone-deprived media. Luciferase activity, indicative of peptide-ERα interaction, was measured and normalized to β-galactosidase activity to provide a quantitative measure of peptide-ERα binding in a cellular context under different ligand conditions.

### Luminescence measurements of E2-mediated ER-SRC3 interaction

HEK293 cells were seeded at 12,000 cells/well in white 96-well plates with 100 μL DMEM containing 8.5% charcoal-stripped FBS. Cells were incubated overnight (37°C, 5% CO2, humidified atmosphere). The following day, cells were transfected with plasmids encoding ER and SRC3 fused to NanoLuc luciferase fragments using FuGENE HD transfection reagent (Promega, # E2311) at a 3:1 lipid:DNA ratio. After 48 hours, 25 μL of 5X Nano-Glo Luciferase assay substrate (Promega, # N1120) was added per well in Nano-Glo LCS dilution buffer (Promega, #N1150). Following a 5-minute room temperature incubation, baseline luminescence was measured using a FLUOstar Omega microplate reader (BMG LabTech). Cells were treated with vehicle or E2 at specified concentrations, and luminescence was measured continuously for 54 minutes.

### Cell growth assay

Six cell lines were used: MCF7-WT, MCF7-Y537S, MCF7-D538G, T47D-WT, T47D-Y537S, and T47D-D538G. These ER knock-in cell lines were previously generated and validated as described^16,18^. Cells were seeded in 96-well plates at 1.5 ×10^3^ or 3.0 × 10^3^ cells per well. Cells were treated with ethanol vehicle (Thermo Scientific, #T038181000), 4OH-tamoxifen (Cayman, #17308), fulvestrant (Selleckchem, #S1191), and/or MTO (ApexBio #B2114). Treatment media were refreshed every 48 hours. Cell numbers were quantified on days 0, 3, 5, and 7 post-treatments using the Cell Counting Kit-8 (CCK-8; Dojindo, #CK04) according to the manufacturer’s instructions. Absorbance at 450 nm was measured using a SPECTRAMax M2 microplate reader (Molecular Devices). Cell viability was calculated as a percentage relative to the ethanol vehicle control for each time point. Data were analyzed using GraphPad Prism 8 software. Results are presented as mean ± standard deviation (SD) from four independent experiments performed in triplicate.

### Migration and Invasion assays

Transwell inserts (8 µm pore size; Corning, #3428) in 24-well plates were used for migration and invasion assays. For invasion assays, insert membranes were coated with 40 μg/ml type I collagen (Corning, #CLS354231). In both assays, 2×10⁵ cells were seeded in the upper chamber in serum-free medium, with 10% fetal bovine serum (FBS; R&D Systems, #S11150) medium in the lower chamber as a chemoattractant. After 48 hours at 37°C, non-migrated/non-invaded cells were removed from the upper chamber. Migrated or invaded cells on the lower surface were fixed with 4% paraformaldehyde (Millipore Sigma, #158127) for 15 minutes, then stained with 1 μg/ml DAPI (Thermo Scientific, #62248) for 10 minutes at room temperature. Stained cells were counted in three random fields per insert using a fluorescence microscope (OLYMPUS, BX43). Assays were performed in triplicate, with three independent experimental replicates. Results are presented as mean ± standard deviation of cells per field.

### Soft agar colony formation assay

Cells (2000 per well) were seeded in duplicate in 6-well plates containing 0.3% top agar over a 0.6% base agar layer. Cultures were maintained at 37°C, 5% CO2 for 14 days, with 100 μL of fresh medium added twice weekly. Colonies were then fixed with 4% paraformaldehyde (15 minutes) and stained with 0.1% crystal violet (10 minutes) at room temperature. After gentle PBS washing and air-drying, standardized digital images were captured. Colony quantification (diameter >50 μm) was performed using ImageJ (version 1.53; NIH). Results represent mean colony count ± standard deviation from three independent experiments.

### 3D spheroid growth assay

MCF7 cells were cultured as 3D spheroids in 96-well microplates (Corning, #3598) precoated with 50 μL of 1.5% agarose. After agarose solidification, 3000 cells in 200 μL complete growth medium were seeded per well and incubated for 24 hours to form single spheroids. Spheroids were treated with 0.1% DMSO vehicle, 1 μM 4OH-tamoxifen, 1 μM fulvestrant, or 1 μM MTO. Every 48 hours, 100 μL of the medium was carefully replaced with a fresh treatment-containing medium to maintain spheroid integrity and constant ligand concentrations. Spheroid growth was monitored by imaging at 48-hour intervals using an ECLIPSE TS2R microscope (Nikon). Spheroid diameter and area were quantified using ImageJ (version 1.53; NIH). The experiment was performed in technical triplicate and repeated three times independently. This protocol was adapted from the literature^58^.

### Orthotopic xenografts for in vivo animal studies

NOD-SCID female mice were maintained under specific pathogen-free conditions in the Athymic Animal Core Facility at the Case Comprehensive Cancer Center. At 5-6 weeks of age, mice underwent orthotopic mammary fat pad injection of cells. MCF7-Y537 cells (1×10^6^) were suspended in PBS and mixed 1:1 with matrigel (R&D Systems, #343300501). The cell-matrigel mixture was homogenized before implantation. A vertical incision was made in the skin between the fourth pair of mammary glands, avoiding damage to the peritoneum. The fourth mammary gland was dissected to expose the mammary fat pad. Using a syringe, 100 μL of the cell-matrigel mixture was injected bilaterally into the mammary fat pads. A 17β-estradiol pellet (1.7 mg, 60-day release; Innovative Research of America, #SE-121) was implanted subcutaneously. Tumor dimensions were measured using digital calipers. Tumor volume was calculated as W^2^×L/2, where W is width and L is length. When tumor volume reached approximately 62.5 mm^3^ (5 mm × 5 mm), mice were randomized into four treatment groups: 1) vehicle (PBS), 2) fulvestrant (50 mg/kg, subcutaneously, twice/week), 3) MTO (1 mg/kg, intraperitoneal, once/week), and 4) combination (fulvestrant 30 mg/kg, subcutaneously, twice/week; MTO 0.5 mg/kg, intraperitoneal, once/week). After 30 days of treatment, all mice were humanely euthanized, and tumors were harvested for further analysis.

### Western blot and protein turnover analysis

Cells were lysed using RIPA buffer (150 mM NaCl, 1% Nonidet P-40, 0.5% DOC, 0.1% SDS, 50 mM Tris-HCl, pH 7.4) supplemented with protease inhibitors (Roche, #11697498001). Proteins (30 μg per lane) were separated on 10% SDS-PAGE gels and transferred to PVDF membranes (Bio-Rad, #1620177). Membranes were blocked with 5% non-fat milk in PBST for 1 hour at room temperature, then incubated with primary antibodies overnight at 4°C. Primary antibodies included: β-actin (1:5000, Cell Signaling, #4967), Lamin B1 (1:1000, Santa Cruz, #sc-374015), α-tubulin (1:2000, Santa Cruz, #sc-166729), HA-tag (1:1000, Santa Cruz, #sc-7392), ER (1:1000, Abcam, #ab16660), Chk1 (1:1000, Cell Signaling, #2360), p-Chk1 (1:1000, Cell Signaling, #2348), Chk2 (1:1000, Cell Signaling, #2662), and p-Chk2 (1:1000, Cell Signaling, #2661). After washing, membranes were incubated with HRP-conjugated secondary antibodies (1:5000, Bio-Rad, #1706515 or #1706516) for 1 hour at room temperature. Protein bands were visualized using an ECL substrate (Thermo Scientific, #32132) and imaged on a ChemiDoc MP system (Bio-Rad, #12003154). Where indicated, cells were treated with fulvestrant (1 μM, Selleckchem, #S1191), MTO (1 mM, ApexBio #B2114), VE-821 (1 μM, Selleckchem, #S8007), KU-55933 (10 μM, Selleckchem, #1092), cycloheximide (20 μM, Millipore Sigma, #66-81-9), or MG132 (10 μM, Selleckchem, #S2619) for 24 hours before lysis. Band intensities were quantified using Fiji ImageJ and normalized to loading controls. Data were analyzed and visualized using GraphPad Prism 8, with results presented as mean ± SD from 3 independent experiments.

To assess protein turnover rates, equal numbers of cells were seeded into 6-cm dishes and cultured for 24 hours. Cells were then pre-treated with cycloheximide (20 μM) for 30 minutes to inhibit new protein synthesis, followed by treatment with DMSO (vehicle control) or MTO (1 μM). Cells were harvested by centrifugation at 0-, 2-, 4-, and 8-hours post-treatment. For proteasome inhibition experiments, cells were pre-incubated with MG132 (10 μM) for 30 minutes, followed by treatment with DMSO, fulvestrant (1 μM), MTO (1 μM), or doxorubicin (DOX, 1 μM) for 8 hours. Cell pellets were lysed in an appropriate lysis buffer, and protein levels were analyzed by western blotting, using β-actin as a loading control. Protein levels were quantified by densitometry and normalized for loading control. The half-life of target proteins, particularly ER, was calculated based on the decay curve fitted to the normalized protein levels over time.

### Immunofluorescence microscopy

MCF7 and T47D cells were cultured in their respective media (DMEM with 10% FBS for MCF7; RPMI with 10% FBS and 0.2 units/ml bovine insulin for T47D) on sterilized round coverslips in 12-well plates. Cells were treated with DMSO (vehicle control) or MTO at specified concentrations and durations. Cells were washed with ice-cold PBS, fixed with 4% paraformaldehyde in PBS for 10 minutes, and permeabilized with 0.1% Triton X-100 in 4% paraformaldehyde for 10 minutes. After six PBS washes, cells were blocked with 3% BSA in PBS for 1 hour, then incubated overnight at 4°C with primary antibody (Abcam, #ab16660; 1:500 in 3% BSA-PBS). Following six PBST washes, cells were incubated with Alexa Fluor 488-conjugated secondary antibody (Invitrogen, #A32731; 1:1000 in PBS) for 1 hour. After six final PBST washes, coverslips were mounted on glass slides using a DAPI-containing medium (Vectashield, #H-2000). Images were captured using an Olympus BX-43 immunofluorescence microscope with a 40× objective under consistent settings.

### In-cell western assay

MCF7 cells (25,000 cells/well) were seeded in 96-well clear-bottom black plates with DMEM/F12 medium containing 8% charcoal-dextran-treated FBS. After 48 hours, cells were treated with hormone at concentrations ranging from 10⁻¹² to 10⁻⁶ M for 24 hours. Cells were then fixed with 3.7% formaldehyde, permeabilized with 0.1% Triton X-100 in PBS, and blocked with In-Cell Western buffer (3% goat serum, 1% BSA, 0.1% cold fish skin gelatin, 0.1% Triton X-100 in PBS). Cells were incubated overnight with anti-ER antibody (Fisher Scientific, # PIMA514501, 1:500) in diluted blocking buffer (1:3 in PBS). After washing with 0.1% Tween in PBS, cells were stained with CF770 goat anti-rabbit secondary antibody (Biotium, # CF770, 1:2000) in a diluted blocking buffer. ER protein expression was quantified using a LI-COR Odyssey imaging system and normalized to DRAQ5 DNA stain (Thermo Scientific, # 62251, 1:10,000). Results are expressed as the percentage of ER remaining after drug treatment compared to the DMSO control.

### Gene expression analysis by RNA-sequencing

Total RNA was extracted from wildtype, Y537S, and D538G ER knock-in MCF-7 cells treated with vehicle control, fulvestrant (1 μM), MTO (1 μM), or combination treatment for 48 hours using RNeasy Plus Mini Kit (Qiagen, #74134). cDNA libraries were generated and sequenced at the on-campus Genomics Core Facility using NextSeq 550 v2.5 for 75 bp single-end reads, with two biological replicates per condition. Reads were processed with FastQC to trim adaptor sequences and aligned to the human genome (GENCODE v38) using HISAT2^59^. Differential expression analyses were performed using DESeq2^60^.

Raw read counts underwent normalization to Reads Per Kilobase Million (RPKM) and subsequent conversion to Transcripts Per Million (TPM), adjusting for sequencing depth and gene length variations. The clusterProfiler package was utilized for Gene Ontology (GO) and Kyoto Encyclopedia of Genes and Genomes (KEGG) pathway enrichment analyses^61^. Gene Set Enrichment Analysis (GSEA) was performed to identify significantly enriched gene sets between experimental conditions^62^. To assess pathway activity variations across samples, Gene Set Variation Analysis (GSVA) was conducted using the GSVA package^63^.

Principal component analysis (PCA) was performed to visualize global gene expression profiles across all conditions, with PC1 and PC2 explaining 31% and 14% of the total variance, respectively. Heatmaps were generated to depict the relative expression patterns of early (4-6 hours) and late (24 hours) estrogen-responsive genes in wild-type and mutant ER cells. The Mfuzz package in R, based on the fuzzy c-means algorithm, was used for soft clustering of gene expression patterns. Z-score normalization was applied to gene expression values, with red indicating upregulation and blue indicating downregulation, compared to the vehicle control group.

## Acknowledgments

We thank Drs. Rinath Jeselsohn and Myles Brown (Dana-Farber Cancer Institute) for providing the MCF7 ER mutant knock-in cell lines, and Drs. Steffi Oesterreich and Adrian Lee (University of Pittsburgh) for the T47D ER mutant knock-in cell lines.

## Author contributions

SY and HYK conceived and designed the study. SY performed computational docking. YL conducted recombinant protein purification and binding measurements. YP performed fluorescence quenching experiments. SY and LG designed umbrella sampling molecular dynamics simulations, executed by QW. HYK and SY designed in-cell experiments executed by HW, ZY, CW, ZL, XL, CP, YC, and JY. SA and SP conducted phage display, dose-dependent reporter assays, in-cell western blots, and SRC3 interaction studies under the guidance of DPM. SY, HW, and HYK designed xenograft studies using NOD/SCID mice, executed by HW and YC. KLW initially conducted toxicity studies using NSG mice under the guidance of RAK. SY and HYK wrote the original draft of the manuscript. All authors contributed to the editing and review of the manuscript and approved the final version.

