## supplemental for "Mitoxantrone inhibits and downregulates ER*α* through binding at the DBD-LBD interface"

**Supplementary Figures**

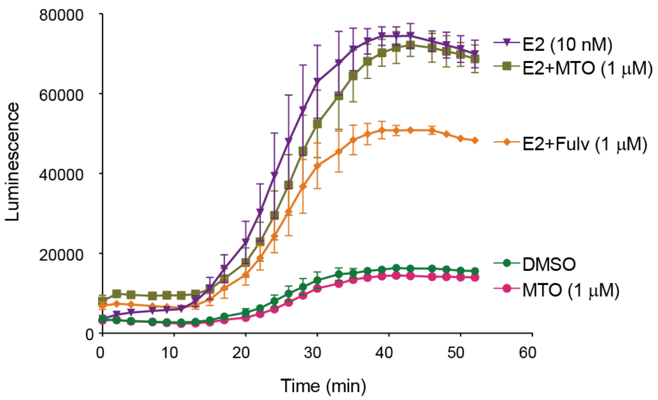


**Figure S1. MTO, unlike fulvestrant, maintains E2-mediated ER-SRC3 interaction.** HEK293 cells were cultured in DMEM with 8.5% charcoal-stripped FBS (12,000 cells/well, 96-well plate) for 24h, then transfected with ER and SRC3 expression plasmids using FuGENE 6 HD. After 48h, protein-protein interactions were assessed using a Nano-Glo Luciferase Assay. Cells were treated with vehicle (DMSO), E2 (10 nM), fulvestrant (1 µM), or MTO (1 µM). Luminescence was measured continuously for 54 min using a FLUOstar Omega plate reader. While fulvestrant disrupted the E2-mediated ER-SRC3 interaction, MTO did not affect it, suggesting a distinct mechanism of action.

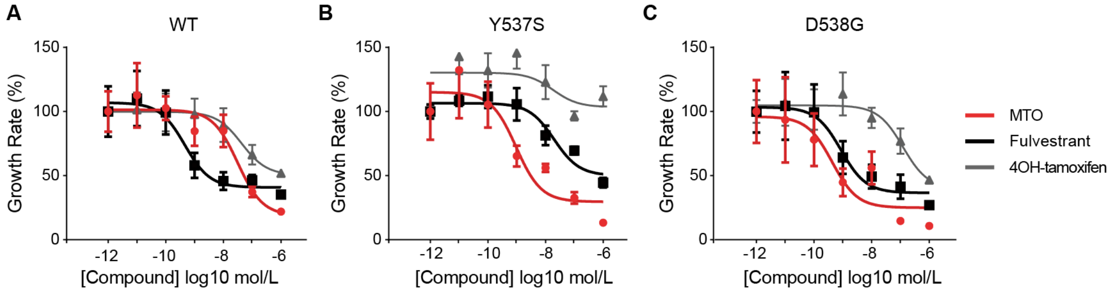


**Figure S2. MTO shows superior anticancer efficacy in endocrine-resistant T47D cells.** MTO exhibited a lower half-maximal inhibitory concentration than 4OH-tamoxifen and fulvestrant in T47D cells harboring ER Y537S and D538G knock-in mutations. Cell viability was assessed over 7 days using a CCK-8 assay, with absorbance measured at 450 nm.

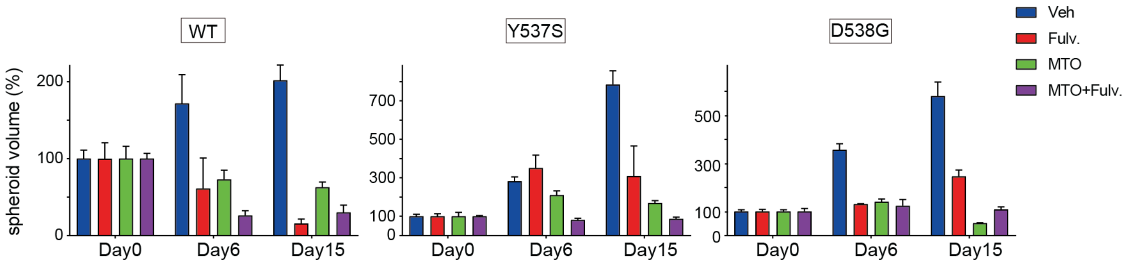


**Fig. S3. Quantification of 3D spheroid growth of MCF7 cells with wild-type and mutant ER.**

Effects of MTO and fulvestrant on MCF7 cells expressing wild-type ER, ER-Y537S, and ER-D538G in 3D culture. Treatments: single agents (1 µM each) or combination (0.5 µM each). Spheroids were imaged bi-weekly using a Nikon ECLIPSE TS2R microscope and quantified with ImageJ (v1.53, NIH). Data shown as mean ± SEM from three independent triplicate experiments. Note: Y-axis scales differ among the three knock-in cell lines. One-way ANOVA with Dunnett's test: *$p < 0.05$; **$p < 0.01$; ***$p < 0.005$.

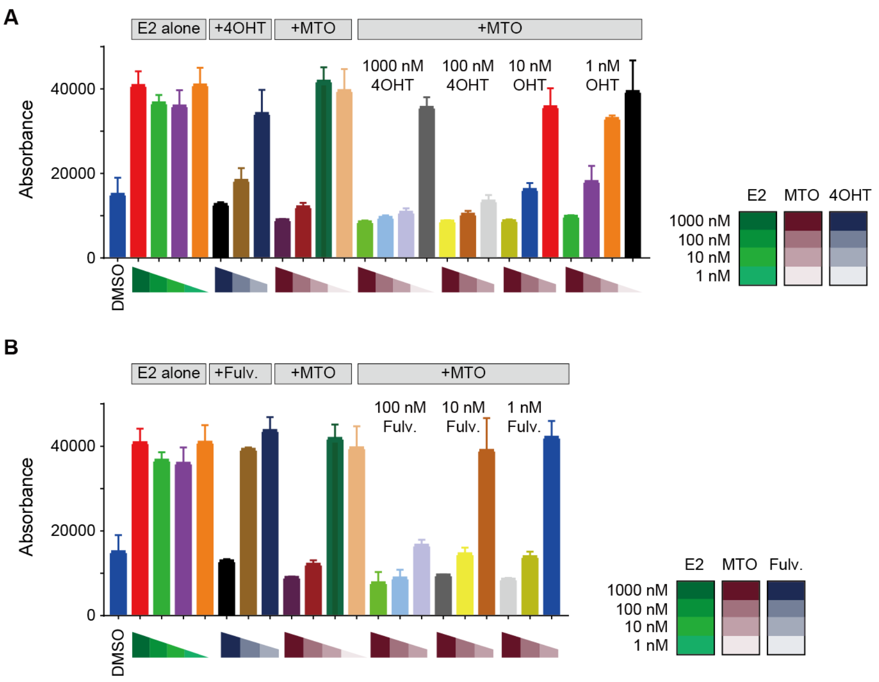


**Figure S4. The effects of MTO in combination with 4OH-tamoxifen or fulvestrant on the growth of MCF7 cells.**

MCF7 cells were plated in DMEM/F12 supplemented with 8% charcoal dextran-treated FBS in 96-well plates (5,000 cells/well) for 48 h. Cells were treated with E2 (dose-response $10^{-9}$-$10^{-6}$ M) (lanes 2-5) with or without ligands (dose-response $10^{-9}$-$10^{-6}$ M) for four days. Plates were decanted and frozen at -80°C overnight before DNA quantitation by fluorescence using Hoechst 33258.

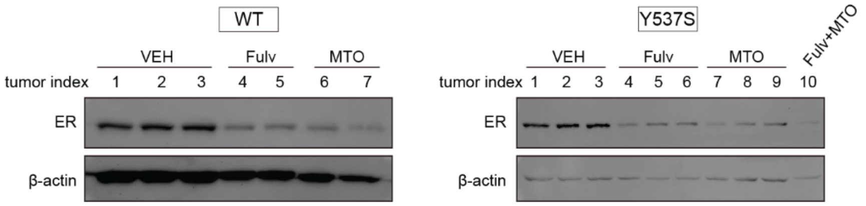


**Figure S5. ER protein levels in MCF7-WT and MCF7-Y537S orthotopic xenografts.** Western blot analysis of ER protein levels in tumors from MCF7-Y537S knock-in orthotopic xenografts in female NOD/SCID mice. Treatment groups: vehicle (VEH), fulvestrant (50 mg/kg, s.c., twice/week), MTO (1 mg/kg, i.p., once/week), and combination (fulvestrant 30 mg/kg, s.c., twice/week + MTO 0.5 mg/kg, i.p., once/week). ER levels were quantified relative to β-actin (loading control). Combination-treated MCF7-WT xenografts were excluded due to insufficient tumor size. Data: mean ± SEM (n = 10 tumors/group).

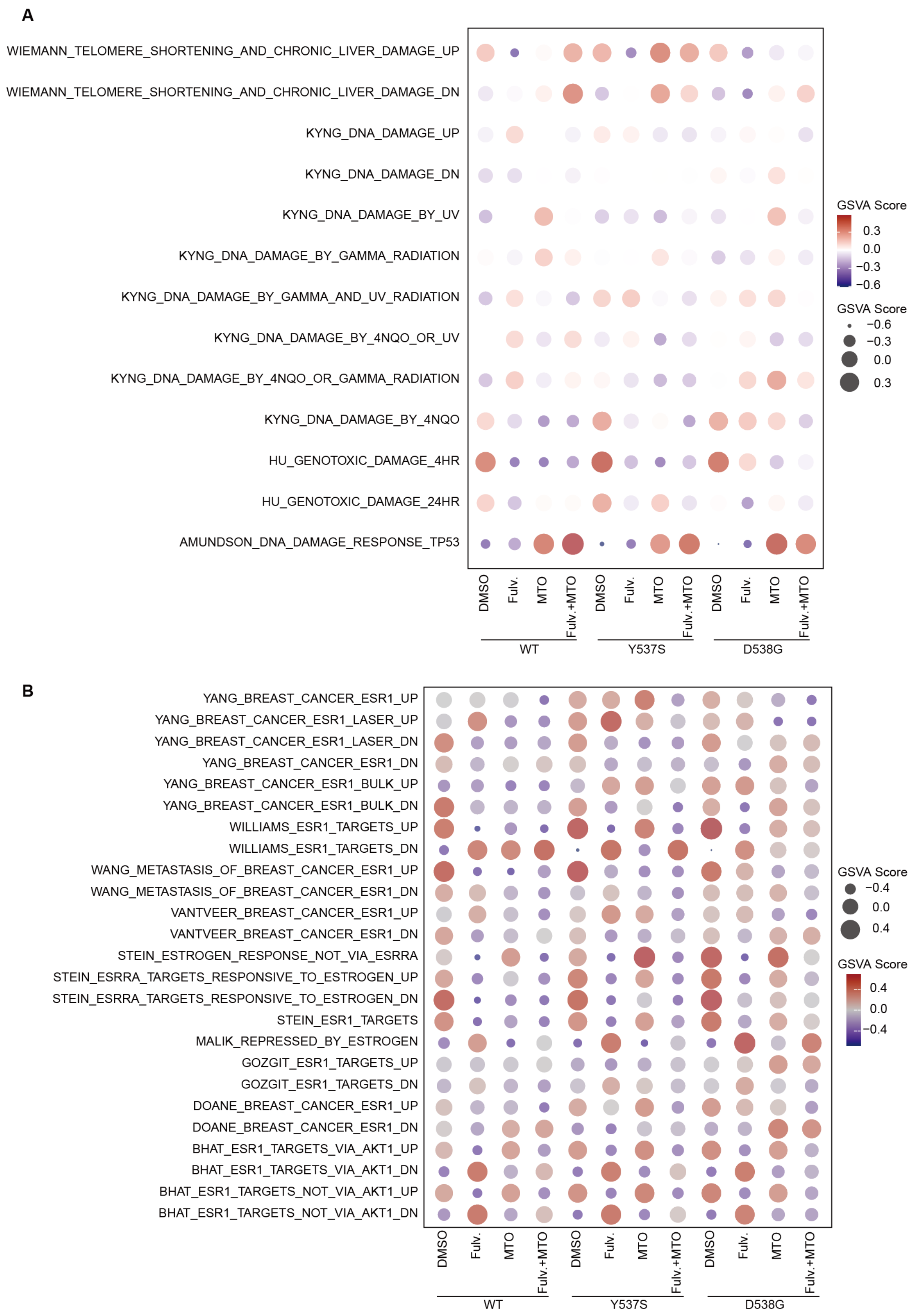
A
WIEMANN_TELOMERE_SHORTENING_AND_CHRONIC_LIVER_DAMAGE_UP
WIEMANN_TELOMERE_SHORTENING_AND_CHRONIC_LIVER_DAMAGE_DN
KYNG_DNA_DAMAGE_UP
KYNG_DNA_DAMAGE_DN
KYNG_DNA_DAMAGE_BY_UV
KYNG_DNA_DAMAGE_BY_GAMMA_RADIATION
KYNG_DNA_DAMAGE_BY_GAMMA_AND_UV_RADIATION
KYNG_DNA_DAMAGE_BY_4NQO_OR_UV
KYNG_DNA_DAMAGE_BY_4NQO_OR_GAMMA_RADIATION
KYNG_DNA_DAMAGE_BY_4NQO
HU_GENOTOXIC_DAMAGE_4HR
HU_GENOTOXIC_DAMAGE_24HR
AMUNDSON_DNA_DAMAGE_RESPONSE_TP53
DMSO
Fulv.
MTO
Fulv.+MTO
WT
Y537S
D538G
GSVA Score
0.3
0.0
−0.3
−0.6
GSVA Score
−0.6
−0.3
0.0
0.3
B
YANG_BREAST_CANCER_ESR1_UP
YANG_BREAST_CANCER_ESR1_LASER_UP
YANG_BREAST_CANCER_ESR1_LASER_DN
YANG_BREAST_CANCER_ESR1_DN
YANG_BREAST_CANCER_ESR1_BULK_UP
YANG_BREAST_CANCER_ESR1_BULK_DN
WILLIAMS_ESR1_TARGETS_UP
WILLIAMS_ESR1_TARGETS_DN
WANG_METASTASIS_OF_BREAST_CANCER_ESR1_UP
WANG_METASTASIS_OF_BREAST_CANCER_ESR1_DN
VANTVEER_BREAST_CANCER_ESR1_UP
VANTVEER_BREAST_CANCER_ESR1_DN
STEIN_ESTROGEN_RESPONSE_NOT_VIA_ESRRA
STEIN_ESRRA_TARGETS_RESPONSIVE_TO_ESTROGEN_UP
STEIN_ESRRA_TARGETS_RESPONSIVE_TO_ESTROGEN_DN
STEIN_ESR1_TARGETS
MALIK_REPRESSED_BY_ESTROGEN
GOZGIT_ESR1_TARGETS_UP
GOZGIT_ESR1_TARGETS_DN
DOANE_BREAST_CANCER_ESR1_UP
DOANE_BREAST_CANCER_ESR1_DN
BHAT_ESR1_TARGETS_VIA_AKT1_UP
BHAT_ESR1_TARGETS_VIA_AKT1_DN
BHAT_ESR1_TARGETS_NOT_VIA_AKT1_UP
BHAT_ESR1_TARGETS_NOT_VIA_AKT1_DN
DMSO
Fulv.
MTO
Fulv.+MTO
WT
Y537S
D538G
GSVA Score
−0.4
0.0
0.4
GSVA Score
0.4
0.0
−0.4

**Figure S6. MTO and fulvestrant modulate gene expression in DNA-damage response and ER signaling pathways.**

Gene Set Variation Analysis (GSVA) of RNA-Seq data using gene sets from the Molecular Signatures Database (MSigDB) C2 Chemical and Genetic Perturbations (CGP) collection[62,64].

(**A**) Heatmap showing MTO and fulvestrant impact on DNA-damage response pathways.

(**B**) Heatmap depicting downregulation of estrogen-related signaling by MTO and fulvestrant. Their combination exhibits the most potent effect.

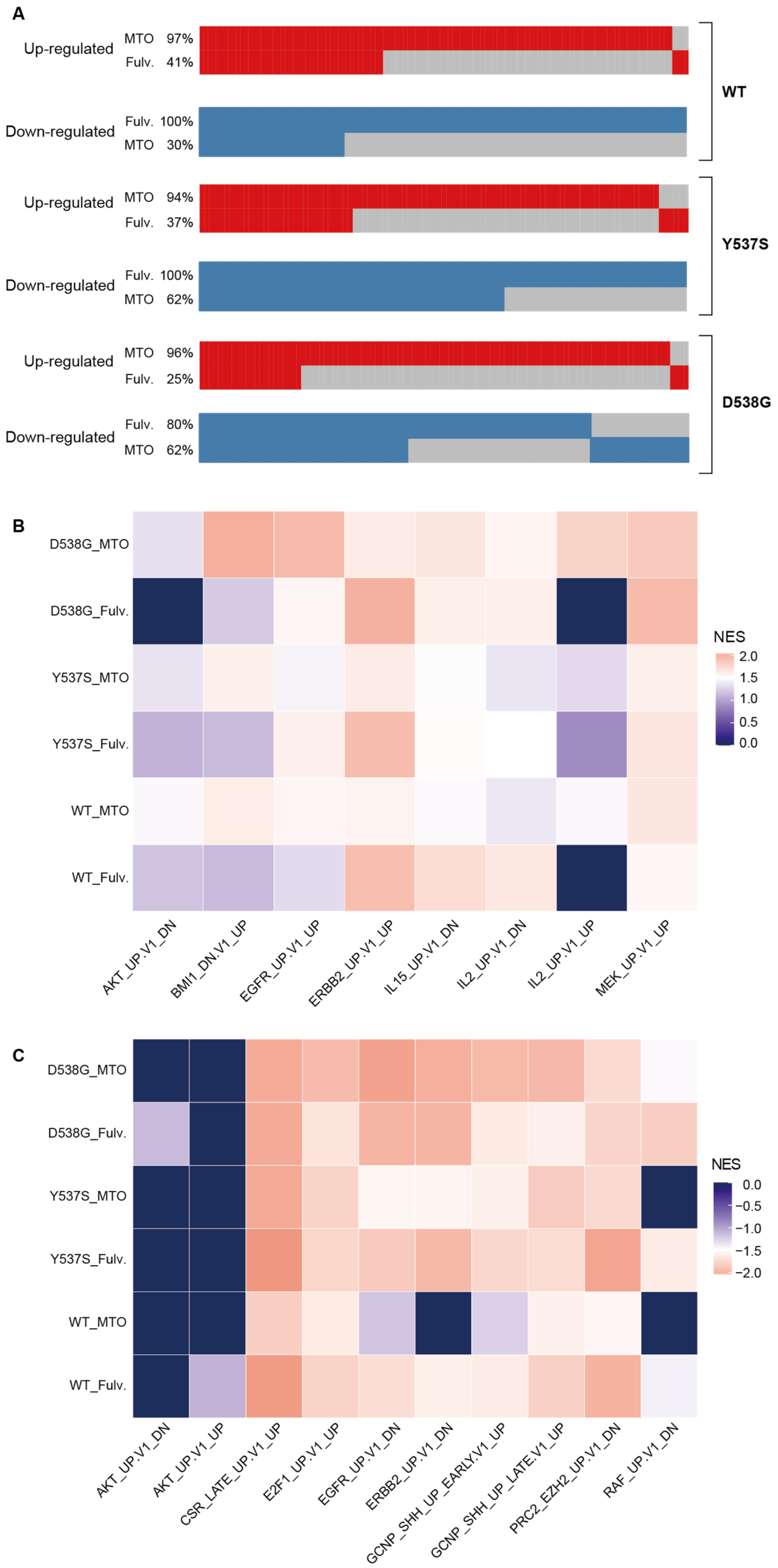
A
Up-regulated
MTO 97%
Fulv. 41%
Down-regulated
Fulv. 100%
MTO 30%
WT
Up-regulated
MTO 94%
Fulv. 37%
Down-regulated
Fulv. 100%
MTO 62%
Y537S
Up-regulated
MTO 96%
Fulv. 25%
Down-regulated
Fulv. 80%
MTO 62%
D538G
B
D538G_MTO
D538G_Fulv.
Y537S_MTO
Y537S_Fulv.
WT_MTO
WT_Fulv.
NES
2.0
1.5
1.0
0.5
0.0
AKT_UP.V1_DN
BMI1_DN.V1_UP
EGFR_UP.V1_UP
ERBB2_UP.V1_UP
IL15_UP.V1_DN
IL2_UP.V1_DN
IL2_UP.V1_UP
MEK_UP.V1_UP
C
D538G_MTO
D538G_Fulv.
Y537S_MTO
Y537S_Fulv.
WT_MTO
WT_Fulv.
NES
0.0
−0.5
−1.0
−1.5
−2.0
AKT_UP.V1_DN
AKT_UP.V1_UP
CSR_LATE_UP.V1_UP
E2F1_UP.V1_UP
EGFR_UP.V1_DN
ERBB2_UP.V1_DN
GCNP_SHH_UP_EARLY.V1_UP
GCNP_SHH_UP_LATE.V1_UP
PRC2_EZH2_UP.V1_DN
RAF_UP.V1_DN

**Figure S7. MTO and fulvestrant differentially impact gene expression pathways.**
**(A)** Waterfall plots of GSEA results from RNA-Seq data, highlighting distinct effects of MTO and fulvestrant on oncogenic signature gene sets in wildtype and ER mutant (537S and D538G) cells.
(**B**)-(**C**). Heatmaps depicting upregulated (B) and downregulated (C) oncogenic signature gene sets in response to MTO and fulvestrant treatments. Color intensity represents normalized enrichment scores (NES).

**Supplementary Table 1. Top 10 candidates identified from computational docking of the NIH Clinical Collection**

CODENAME: Ranked identifier (T1-T10) based on docking score
PUBCHEM_SID: Unique compound identifier in PubChem database
NCC_SAMPLE_ID: Sample identifier from the NIH Clinical Collection
COMPOUND_NAME: Generic or trade name of the compound

| CODENAME | PUBCHEM_SID | NCC_SAMPLE_ID | COMPOUND_NAME |
|---|---|---|---|
| T1 | 46387005 | SAM001247096 | Salmeterol |
| T2 | 46386998 | SAM001247089 | HTMT dimaleate |
| T3 | 46386763 | SAM001246750 | Aripiprazole |
| T4 | 104170122 | SAM002699896 | Mitoxantrone |
| T5 | 104170163 | SAM002548975 | Raloxifene |
| T6 | 46386937 | SAM001246863 | Fluphenazine |
| T7 | 46386544 | SAM001246526 | Cladribine |
| T8 | 104170169 | SAM002699891 | Racepinephrine |
| T9 | 46386749 | SAM001246736 | Carvedilol |
| T10 | 46386802 | SAM001246975 | Cisapride monohydrate |
